# Threat and reward imminence processing in the human brain

**DOI:** 10.1101/2023.01.20.524987

**Authors:** Dinavahi V. P. S. Murty, Songtao Song, Srinivas Govinda Surampudi, Luiz Pessoa

**Affiliations:** Department of Psychology, University of Maryland, College Park, MD, USA

**Author notes:** Corresponding author: Luiz Pessoa.

## Abstract

In the human brain, aversive and appetitive processing have been studied with controlled stimuli in rather static settings. In addition, the extent to which aversive- and appetitive-related processing engage distinct or overlapping circuits remains poorly understood. Here, we sought to investigate the dynamics of aversive and appetitive processing while male and female participants engaged in comparable trials involving threat-avoidance or reward-seeking. A central goal was to characterize the temporal evolution of responses during periods of *threat or reward imminence*. For example, in the aversive domain, we predicted that the bed nucleus of the stria terminalis (BST), but not the amygdala, would exhibit anticipatory responses given the role of the former in anxious apprehension. We also predicted that the periaqueductal gray (PAG) would exhibit threat-proximity responses based on its involvement in proximal-threat processes, and that the ventral striatum would exhibit threat-imminence responses given its role in threat escape in rodents. Overall, we uncovered imminence-related temporally increasing (“ramping”) responses in multiple brain regions, including the BST, PAG, and ventral striatum, subcortically, and dorsal anterior insula and anterior midcingulate, cortically. Whereas the ventral striatum generated anticipatory responses in the proximity of reward as expected, it also exhibited threat-related imminence responses. In fact, across multiple brain regions, we observed a main effect of arousal. In other words, we uncovered extensive temporally-evolving, imminence-related processing in both the aversive and appetitive domain, suggesting that distributed brain circuits are dynamically engaged during the processing of biologically relevant information irrespective of valence, findings further supported by network analysis.

**Significance Statement:** In the human brain, aversive and appetitive processing have been studied with controlled stimuli in rather static settings. Here, we sought to investigate the dynamics of aversive/appetitive processing while participants engaged in trials involving threat-avoidance or reward-seeking. A central goal was to characterize the temporal evolution of responses during periods of *threat or reward imminence*. We uncovered imminence-related temporally increasing (“ramping”) responses in multiple brain regions, including the bed nucleus of the stria terminalis, periaqueductal gray, and ventral striatum, subcortically, and dorsal anterior insula and anterior midcingulate, cortically. Overall, we uncovered extensive temporally-evolving, imminence-related processing in both the aversive and appetitive domain, suggesting that distributed brain circuits are dynamically engaged during the processing of biologically relevant information irrespective of valence.

## Introduction

In the human brain, aversive and appetitive processing have been studied with controlled stimuli in rather static settings, including emotional faces and classical conditioning in the aversive domain and reward cues in the appetitive domain. In these studies, stimuli are typically short in duration and consequently lack the temporal dynamics encountered in more natural settings (Adolphs et al., 2016). In addition, the extent to which aversive- and appetitive-related processing engage distinct or overlapping circuits remains poorly understood (Leknes and Tracey, 2008; Bissonette et al., 2014; Hayes et al., 2014; Pessiglione and Delgado, 2015).

Studies of sustained threat processing (Somerville et al., 2010, 2013; Alvarez et al., 2011; McMenamin et al., 2014; Meyer et al., 2019b; Hur et al., 2020; Murty et al., 2022) have uncovered anticipatory responses to threat. For example, Hur et al. (2020) reported increased responses during anticipation of uncertain versus certain threat across multiple brain regions, including the midcingulate cortex, anterior insula, periaqueductal gray, and bed nucleus of the stria terminalis (BST). Mobbs et al. (2007) provided evidence that the periaqueductal gray (PAG) increases responses when threat is proximal (see also Mobbs et al., 2010). In the appetitive domain, anticipatory responses have been observed in the ventral striatum/nucleus accumbens in humans (Knutson and Greer, 2008) and in rodents (Howe et al., 2013). However, functional MRI studies typically employ canonical modelling of responses (i.e., expected responses are obtained via convolution with a hemodynamic filter). Such approach assumes that sustained responses are constant throughout periods of extended threat (Alvarez et al., 2011; Somerville et al., 2013; Hur et al., 2020), and thus do not capture the potential temporal evolution of threat/reward processing. Thus, despite progress, we still lack understanding of the unfolding of aversive and appetitive signals during temporally extended periods.

To address existing gaps in the literature, in the present experiment, we sought to investigate the dynamics of aversive and appetitive processing while participants engaged in comparable trials involving threat-avoidance or reward-seeking. A central goal was to characterize the temporal progression of responses during periods of *threat or reward imminence* (Figure 1). We use the term “imminence” based on an influential model of predatory imminence proposed by Fanselow and Lester (1988; Fanselow, 1994). Predatory imminence is linked to a prey’s assessment of the level of threat posed by a predator and is central to determining defensive behaviors in ecological settings. According to the framework, defensive behaviors depend on a continuum of predator-prey relationships. For example, defensive behaviors circa-strike may be quite dramatic, as the prey seeks to elude the predator just before capture, and differs considerably from defensive behaviors before contact is more imminent. For related work, see Blanchard and Blanchard (1988); McNaughton and Corr (2004); Blanchard et al. (2011) and Mobbs et al. (2015). Given that in our design threat and reward trials had similar temporal properties, we refer to “imminence” in both contexts.

**Figure 1.**
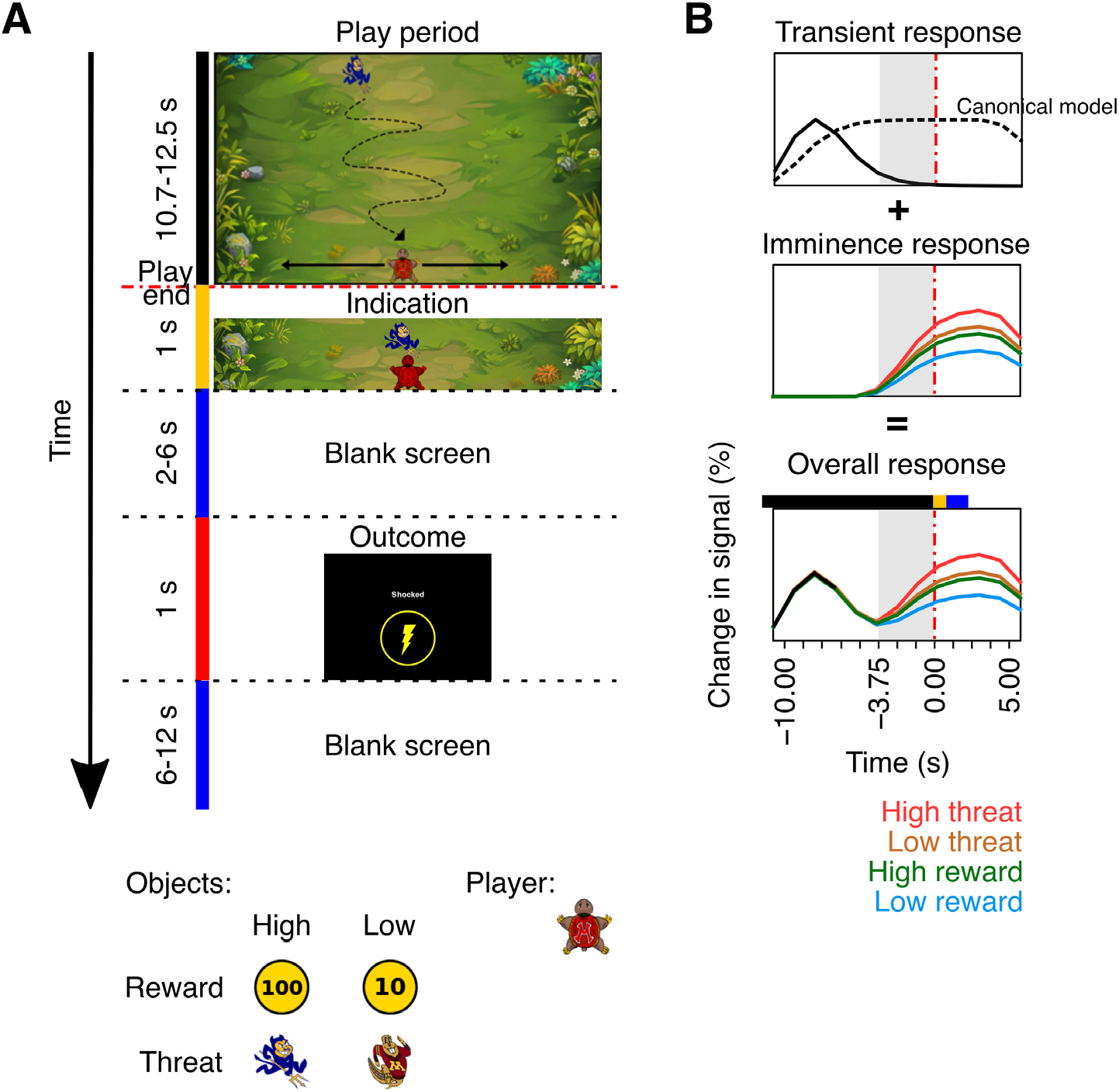
Experimental design. (A) The temporal evolution is indicated vertically. At the outset of the trial, an icon appeared at the top of the screen and descended its length in around 12 seconds. The participant controlled the turtle at the bottom which could only move horizontally. During the indication period, the turtle icon either turned red (if caught by the threat) or green (if it caught the reward) or did not change color (if the subject escaped the threat or missed the reward). The inset shows the icons used as a function of experimental condition, and the icon controlled by the participant (“player”). (B) Schematic responses illustrating our primary hypothesis: potential imminence-related responses. Here, responses assumed a brief initial transient response followed by imminence-related responses for different threat and reward levels. The final simulated response was obtained by summing transient and sustained hypothesized components. The gray zone indicates the temporal window considered for analysis at trial end. The dashed line in the top row represents a canonical hemodynamic response filter convolved with boxcar lasting for the duration of the stimulus, as was assumed in most prior studies, for comparison.

In our paradigm, during threat trials a virtual predator descended across the screen, and the player had to avoid being caught to prevent the delivery of an unpleasant shock (“high”) or a benign electrical stimulation (“low”). During reward trials, a coin likewise descended across the screen, and the player had to catch it to obtain the cash reward (“low” and “high” values). We hypothesized that in such avoidance and approach conditions, we would observe the temporal progression of brain signals as threat/reward imminence increased in space and time, which we call imminence-related responses. Figure 1B illustrates potential imminence-related responses for different threat and reward levels. We estimated these responses without assuming a canonical hemodynamic filter, and statistically evaluated the contributions of valence and arousal within a temporal window near the trial end (−3.75 to 0 seconds of trial end).

One of the goals of our study was to test key predictions related to specific brain regions. First, we hypothesized that increasing imminence-related responses would be observed in brain areas such as the PAG (Fanselow, 1994; Mobbs et al., 2015; Branco and Redgrave, 2020) and the BST (Davis et al., 2010; Tovote et al., 2015), but not the amygdala. Second, we expected that the ventral striatum would exhibit anticipatory responses during reward approach (Knutson and Greer, 2008), as well as during aversive trials given the region’s proposed involvement in threat escape (LeDoux et al., 2017). Third, studies of sustained/uncertain threat have observed responses in the anterior insula but have not clearly separated the contributions of dorsal and ventral parts of the region; e.g. (Alvarez et al., 2011; Grupe et al., 2013; Hur et al., 2020). Here, we hypothesized that the dorsal but not the ventral anterior insula would exhibit threat-related imminence-related responses (Murty et al., 2022). Likewise, we predicted that the anterior MCC but not the ACC would exhibit threat-related imminence responses (Lima Portugal et al., 2020).

Although our main focus was to test the effect of arousal and valence for a set of regions of interest (ROIs), we also interrogated our data at the voxel level to test for the presence of these effects throughout the brain. Finally, we explored the possibility that the regions that showed similar activation profiles also co-activated at the trial level, consistent with the notion that they formed functional clusters. Accordingly, we performed network analysis to identify potential clusters.

## Methods

### Subjects

A total of 96 subjects (38 females) were recruited from the University of Maryland, College Park, community. All had normal or corrected-to-normal vision, and reported no neurological disease or current use of psychoactive drug. They provided written informed consent before participating in the study and were paid immediately after the experiment. The study was approved by the University of Maryland, College Park, Institutional Review Board. Out of the 96 subjects, we used data from 11 to perform exploratory analyses, such as defining the anticipatory temporal window; their data were not used further to avoid circularity in our analyses. Data from 5 other subjects were discarded due to excessive head motion (see below). Thus, results are reported for 80 subjects (32 females) aged 21.1 ± 2.8 years (mean ± SD; range: 18-33 years).

### Behavioral task

Each trial began with a “play period” that consisted of either a threat-avoidance or a reward-pursuit task (Figure 1). Subjects controlled a turtle icon (“player”) at the bottom of the screen, which they could move only horizontally (left or right) using two buttons with their right hand (index and middle fingers). Their objective was to actively avoid a threatening object (“threat”) or pursue a rewarding object (“reward”) that smoothly moved towards the bottom of the screen with a constant speed. The object descended with a component of random motion while at the same time tracking the player’s position such that it tended to move towards the player if it was a threat, and away from the player if it was a reward. As the threat/reward object descended, in the last third of the screen, its random movement component considerably decreased, such that it more directly chased the player during threat, or more directly avoided the player during reward. In this manner, the success/failure of a given trial was only ascertained (or apparent) to the participant at trial end, or possibly a fraction of a second prior to trial end. In this manner, our design avoided the possibility that participants felt an early relief in (presumably) avoiding the predator at earlier stages of trial progression; or, relatedly, experiencing early excitement at the (presumed) impending capture of a coin during a reward trial.

Threat and reward trials were of two intensity levels, low and high. For threat trials, the intensity corresponded to different shock levels; for reward trials, the intensity corresponded to different reward levels (participants were informed that they would win cash based on their performance in a random subset of trials; in reality, all participants received the same total remuneration). All four trial types were associated with a different descending icon, such that the subject was aware of the condition type from the outset of the play period.

The play period ended when the icon reached the bottom of the screen or touched the player. For high threat, the play period lasted 10.74-12.44 seconds (mean: 12.13, SD: 0.50); for low threat: 10.72-12.44 seconds (mean: 12.17, SD: 0.48); for high reward: 10.72-12.44 seconds (mean: 11.73, SD: 0.64), and for low reward: 10.72-12.44 seconds (mean: 11.72, SD: 0.63). The play period was followed by an “indication period” (1 second), during which the turtle icon either turned red (if caught by the threat) or green (if it caught the reward) or did not change color (if the subject escaped the threat or missed the reward). This was followed by an inter-stimulus interval (blank screen) that lasted 2-6 seconds.

The play period was followed by an “outcome phase” lasting 1 second. The subject received a highly unpleasant (but not painful) electrical stimulation if they were caught during a high-threat trial, or a very mild (benign, not unpleasant) electrical stimulation if they were caught during a low-threat trial. Electrical stimulation was delivered to the fourth and fifth fingers of the left hand via MRI-compatible electrodes using an electric stimulator (STMISOC connected to STM100C, BIOPAC Systems, Inc, Santa Barbara, CA), accompanied by the words “Shock” and “Touch” (high and low, respectively). During reward trials, the display showed “Rewarded 100” or “Rewarded 10” to indicate reward level. Lastly, the outcome period was followed by a blank screen lasting 2-6 seconds before the next trial started.

We determined the benign and unpleasant electrical stimulation levels for each subject separately for each run to prevent habituation. To determine the benign level, we started with the minimal stimulation level (1 V setting) and increased it until the subject reliably felt minimal stimulation (range: 5-50 V across subjects; mean: 23.17, SD: 9.98 V); 22 subjects recalibrated this level during the experiment). To determine the unpleasant level, we started with a higher stimulation level (10 V setting) and increased it until the subject reported it as “highly unpleasant but not painful” (range: 10-100 V across subjects; mean: 58.92, SD: 19.34 V); 41 subjects recalibrated this level during the experiment). At the end of each run, subjects were asked verbally to rate the electrical stimulations on a 11-point Likert scale: 0 being not unpleasant at all and 10 being painful. Subjects reported a rating of 2.50 ± 1.02 (mean ± SD across 79 subjects; ratings for 1 subject could not be recorded) for the benign level and 6.53 ± 1.02 for the unpleasant level.

All subjects underwent a training session to get familiarized with the task while structural scans were obtained. During the experiment, subjects avoided high threat in 65.9 ± 11.7% trials and avoided low threat in 67.5 ± 12.0% trials (two-sided paired t-test, *t*(79) = -1.21, p=0.23). Similarly, they successfully pursued high reward in 68.8 ± 8.5% trials and low reward in 66.9 ± 8.6% trials (two-sided paired t-test, *t*(79) = 1.57, p=0.12). These behavioral results indicate that subjects pursued reward and avoided threat irrespective of intensity level.

### MRI data acquisition and preprocessing

We collected MRI data using a 3T Siemens TRIO scanner (Siemens Medical Systems) with a 32-channel head coil. We acquired a high resolution T1-weighted MPRAGE anatomical scan (TR: 2400 ms, TE: 2.01 ms, FOV: 256 mm, voxel size: 0.8 mm isotropic), followed by functional echo planar images using a multiband scanning sequence (TR = 1250 ms, TE = 39.4 ms, FOV = 210 mm, and multiband factor of 6). Each image (volume) contained 66 non-overlapping oblique slices oriented 30 degrees clockwise relative to the AC-PC axis. Thus, voxels were 2.2 mm isotropic. We obtained 410 such volumes for each run. We filled the gap between the last volume of the last trial in the run and the end of the run using random pictures of animals/scenes so that every run had 410 volumes (data not analyzed). Thus each run was 512.5 seconds long in duration. We also acquired double-echo field maps (TE = 73.0 ms) with the acquisition parameters matched to the functional data.

Functional images were preprocessed as described in our previous work by using a combination of fMRI packages and in-house scripts (Limbachia et al., 2021; Murty et al., 2022). We aligned the onset times of each slice in a volume to the first acquired slice (slice-timing correction) with Fourier interpolation, using the 3dTshift program of Analysis of Functional Neuroimages (AFNI; Cox, 1996). The first volume of the resultant data was used as a reference volume to correct the rest of the volumes for head motion using AFNI’s 3dvolreg program. This step also generated motion parameters across time for each run which were later used for rejecting runs based on overall head motion artifacts (see below).

To determine whether or not a voxel belonged to the brain (skull stripping), we employed six different fMRI packages (AFNI: Cox, 1996; BrainSuite: Shattuck and Leahy, 2002; FSL: Smith et al., 2004; SPM: Friston et al., 2007; ANTs: Avants et al., 2009; ROBEX: Iglesias et al., 2011) on T1-weighted structural data. If 4 out of 6 packages estimated a voxel to belong to the brain, it was retained, otherwise it was discarded. This sought to improve coregistration between functional and anatomical images. Next, we used ANTs to estimate a nonlinear transformation mapping the skull-stripped anatomical T1-weighted image to the skull-stripped MNI152 template (interpolated to 1-mm isotropic voxels). The nonlinear transformations from coregistration/unwarping and normalization were combined into a single transformation that was applied to map volume-registered functional volumes to standard space (interpolated to 2-mm isotropic voxels).

Signal intensity at each voxel was normalized to a mean value of 100 separately for each run. For voxelwise analysis (but not ROI level), data were spatially smoothed with a Gaussian filter (4 mm full-width half-maximum) restricted to gray matter voxels before normalizing the signal intensity.

### Data rejection based on head motion

We excluded runs and subjects with heavier head motion as follows. We estimated the Framewise Displacement (FWD; Power et al., 2014) at every time-point for every run from the motion parameters obtained during preprocessing. First, we excluded runs that had FWD of 4.4 mm (2 voxel lengths) or more at any time-point, and runs that had 25% or more of all time-points with FWD over 1.1 mm. In this manner, we rejected 32/650 runs (4.9%) across 85 subjects from analysis. Finally, we rejected those subjects who had less than 4 acceptable runs, leading to a further rejection of 9 runs (1.4%); a total of 5 subjects were excluded. Thus, we used 609 runs across 80 subjects (mean ± SD: 7.61 ± 0.79; range: 4-8 runs per subject), with a total of 9739 trials across them (mean ± SD: 15.99 ± 0.20; range: 11-16 trials per run). Subjects had 30.4 ± 3.1 trials (minimum of 16 trials) for each of four experimental conditions.

To further reduce the contribution of head motion, we used FSL’s Independent Component Analysis, Automatic Removal of Motion Artifacts (ICA-AROMA) toolbox (Pruim et al., 2015) and fsl_regfilt, and regressed out the components classified as head motion from the data. The runs were then concatenated for each subject.

Potential contributions of white matter and cerebrospinal fluid were minimized by excluding voxels that intersected with masks of the respective tissue types. Finally, we excluded voxels if their mean across time was outside the range of 95-105 (recall that signal intensity was normalized to 100 during preprocessing), or if the standard deviation exceeded 25 (mean ± SD: 7.1 ± 1.6%). This step sought to exclude voxels with relatively poor temporal signal-to-noise ratio (ratio of mean signal to standard deviation across time) at the level of individuals.

### Subject-level analysis

Responses were estimated without assuming the canonical shape of the hemodynamic response. Instead, we used a series of cubic splines as regressors to estimate signals. The design matrix for every subject was defined using AFNI’s *3dDeconvolve* program and fit to the data using the 3dREMLfit program. For estimating trial-averaged responses for each condition, we aligned trials to the end of the play period and modeled responses from -10 to +5 seconds relative to the play end (13 cubic splines given the TR of 1.25 seconds), separately for each condition. The outcome period was modelled by convolving a 1-second square pulse with the canonical hemodynamic responses (gamma variate peaking at 4.7 seconds with a full-width at half maximum of 3.77 seconds by using default parameters p=8.6 and q=0.547 in the *3dDeconvolve* program). Using convolved regressors for the outcome period contributed to keeping regressor collinearity at relatively low levels: the maximum variance inflation factor for the design matrices was 1.56 ± 0.04 (range: 1.46-1.70). Further, to minimize the contribution of larger head movements, we censored the time-points that had the Euclidean norm of the derivatives of the motion parameters larger than 1.1 mm (half the voxel dimension). Finally, we included in the model the six head motion parameters (3 translational and 3 rotational) and their temporal derivatives as nuisance regressors, in addition to linear and nonlinear polynomial terms (up to fourth degree) to account for baseline and slow signal drift.

We performed the above analyses for specific regions of interest (ROIs) as well as voxelwise for the whole brain. We examined responses from a total of 36 structurally and/or functionally defined cortical and subcortical ROIs (18 per hemisphere), which have been suggested to be key regions involved in aversive and appetitive processing in the literature. ROIs were non-overlapping and defined *a priori* based on previous literature as mentioned in Table 1. Preprocessed time-series (without spatial smoothing) data were averaged across all voxels within the ROI before running the model.

**Table 1.**
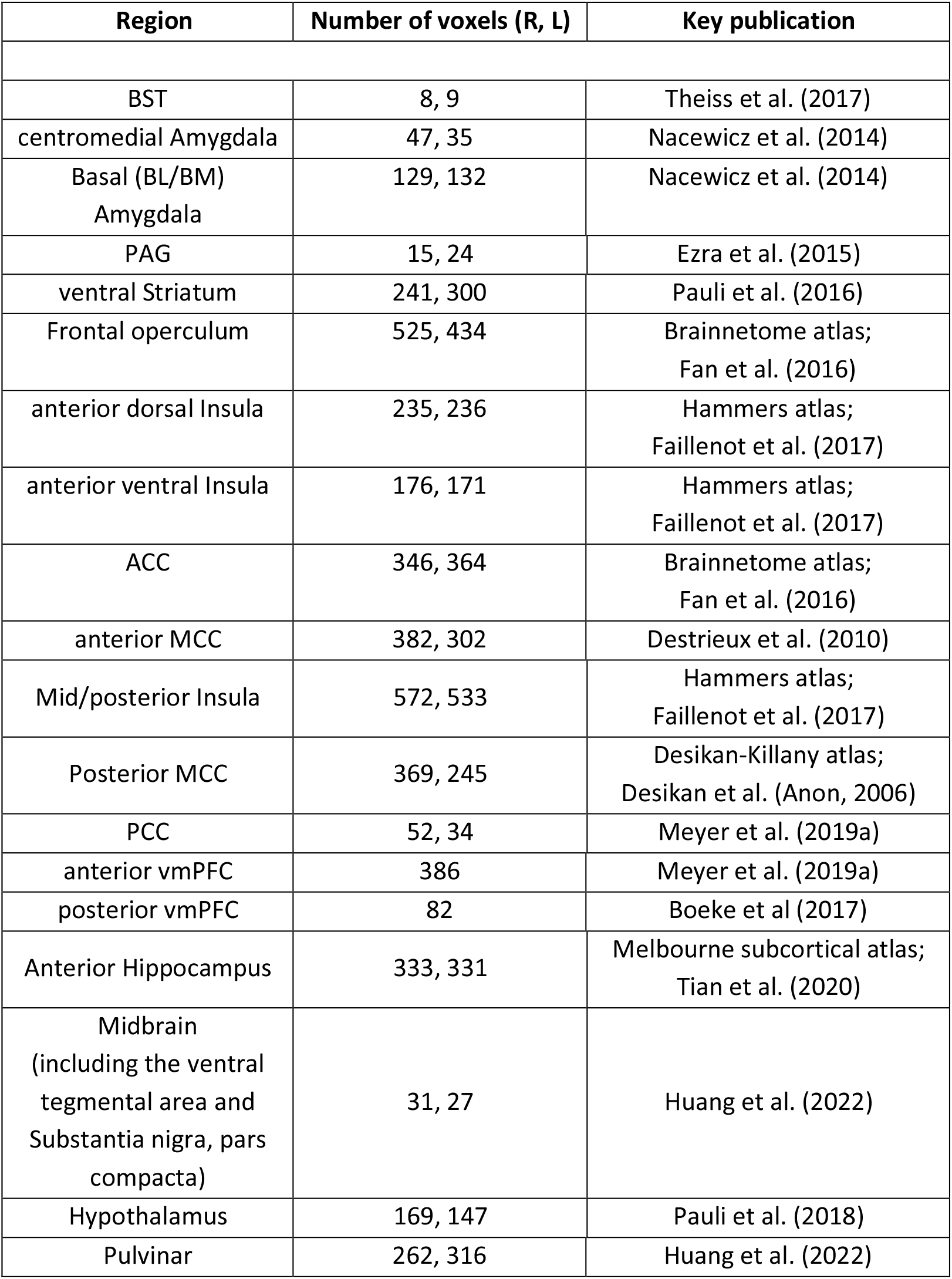
Regions of interest. R, right; L, left.

| Region | Number of voxels (R, L) | Key publication |
| --- | --- | --- |
| BST | 8, 9 | Theiss et al. (2017) |
| centromedial Amygdala | 47, 35 | Nacewicz et al. (2014) |
| Basal (BL/BM)<br>Amygdala | 129, 132 | Nacewicz et al. (2014) |
| PAG | 15, 24 | Ezra et al. (2015) |
| ventral Striatum | 241, 300 | Pauli et al. (2016) |
| Frontal operculum | 525, 434 | Brainnetome atlas;<br>Fan et al. (2016) |
| anterior dorsal Insula | 235, 236 | Hammers atlas;<br>Faillenot et al. (2017) |
| anterior ventral Insula | 176, 171 | Hammers atlas;<br>Faillenot et al. (2017) |
| ACC | 346, 364 | Brainnetome atlas;<br>Fan et al. (2016) |
| anterior MCC | 382, 302 | Destrieux et al. (2010) |
| Mid/posterior Insula | 572, 533 | Hammers atlas;<br>Faillenot et al. (2017) |
| Posterior MCC | 369, 245 | Desikan-Killany atlas;<br>Desikan et al. (Anon, 2006) |
| PCC | 52, 34 | Meyer et al. (2019a) |
| anterior vmPFC | 386 | Meyer et al. (2019a) |
| posterior vmPFC | 82 | Boeke et al (2017) |
| Anterior Hippocampus | 333, 331 | Melbourne subcortical atlas;<br>Tian et al. (2020) |
| Midbrain<br>(including the ventral<br>tegmental area and<br>Substantia nigra, pars<br>compacta) | 31, 27 | Huang et al. (2022) |
| Hypothalamus | 169, 147 | Pauli et al. (2018) |
| Pulvinar | 262, 316 | Huang et al. (2022) |

Estimated responses for fMRI data and skin conductance responses (see below) were plotted by using error bars that take into account within-subject error according to the method by Cousineau and O’Brien (2014).

Finally, note that we were not able to analyze our data as a function of performance (successful vs. unsuccessful trials) given insufficient number of trials across all trial types. As stated, subjects were successful around 66-68% of the time, so only around 30% unsuccessful trials were available on average. Given that participants performed around 30 trials total per condition (both successful and unsuccessful), this left too few repetitions of certain trial types (frequently less than 10 repetitions) to analyze our results in terms of performance.

### Group-level statistical analyses

We conducted group-level analyses on subject-level responses. We defined a temporal window targeting the period towards the end of the play period (−3.75 to 0 seconds), which minimized contributions of potentially confounding processes (e.g., performance-based relief/disappointment at the end of the trial).

The response magnitude for a given trial type was the average of the signals estimated within the temporal window. Such responses were then submitted to a 2 valence (threat, reward) by 2 arousal (low, high) repeated-measures ANOVA, to estimate main effects and the interaction between factors, both at both ROI and voxel levels. Additionally at the ROI level, we tested whether the effects of valence and/or arousal depended on ROI by including ROI as a factor in tests focusing on 1) BST, centromedial and basolateral amygdala; 2) dorsal anterior insula and ventral anterior insula; and 3) ACC and anterior MCC. Thus, we performed 3-way repeated-measures ANOVAs with ROI, valence, and arousal as factors (separately for right and left hemispheres). For ROI data we used the R suite and for voxelwise data we used AFNI’s 3dttest++ to allow censuring of voxels with poor SNR at the subject level.

We corrected for multiple comparisons for ROI-level analyses using False Discovery Rate (false discovery rate of 0.05). For the 2 × 2 analysis, we corrected for a total of 108 tests: 36 ROIs, 2 main effects and 1 interaction. For the 3-way ANOVA, we adjusted the significance level by a factor of 42 (three main effects plus three 2-way interactions plus one 3-way interaction, across 3 groups, and 2 hemispheres; thus, 42 tests in all). Thus, we reported p-values tested against a Bonferroni-corrected significance level of 0.001. For the voxelwise analysis, we applied FDR correction at the level of 0.001 using Python’s scipy.stats package. We corrected based on 156451 (number of voxels) times 3 (2 main effects and 1 interaction effect) tests.

### Skin conductance response acquisition and subject-level analysis

Skin conductance responses were collected using the MP-150 data acquisition system with the GSR100C module (BIOPAC Systems, Inc., Goleta, CA). Signals were acquired at 250 Hz using MRI-compatible electrodes attached to the index and middle fingers of the subjects’ non-dominant, left hand. These signals were then resampled offline to 0.8 Hz (corresponding to a sampling rate of 1.25 s to match the sampling rate of functional MRI data). We used the MATLAB program *resample*.*m* for resampling and used the default anti-aliasing low pass FIR filter (Kaiser window, shape parameter = 5). We performed subject-level analysis on the resulting data in the same way as mentioned for the functional data (but we did not censor time points based on head motion or use motion parameters and their derivatives as nuisance regressors in the model). We conducted group-level analysis following the same approach for fMRI ROIs. In particular, because the time course of skin conductance is very similar to that of the fMRI signal (Gester et al., 2018), we used the same temporal window for analysis.

### Network analysis

Initial network analyses (e.g., testing algorithms, setting parameter values) were performed on a set of 26 participants (including data of 11 pilot subjects; see Subjects). To avoid circularity (“double-dipping”) and enhance the generalizability of our findings, once all processing procedures and parameters were fixed, we applied them to the remaining 65 participants.

For each subject and condition, we estimated responses for each trial separately (using the -*stim_times_IM* option in *3dDeconvolve* of the AFNI package). Note that the maximum variance inflation factor for the design matrices was low (1.36 ± 0.64 (mean ± SD); range: 1.00-4.36). For each subject, we then concatenated ROI-level activations from the analysis window (−3.75 s to 0 s) across trials and computed the Pearson correlation for each pair of ROIs to obtain a functional connectivity matrix, for the high threat and high reward conditions separately. Group-level matrices were then obtained by computing the median functional connectivity matrix across participants, thresholding them (weakest 25% of connections were set to zero), and keeping non-negative weights only (negative entries were set to zero).

Community detection was applied to group-level functional connectivity matrices using the Louvain algorithm (Blondel et al., 2008), which employed a resolution parameter of 1.15 so as to produce approximately 4-6 networks while avoiding breaking them up so much as to result in singletons. Although the Louvain algorithm performs well in practice, it is stochastic in nature, and individual runs of the algorithm may result in different community assignments (Good et al., 2010). To address this issue, we employed consensus clustering (Lancichinetti and Fortunato, 2012; Betzel, 2020) to determine final sets of networks for high threat and for high reward, separately (in each case, consensus was obtained over 1,000 runs of the Louvain algorithm). Finally, we used the pyCircos (Krzywinski et al., 2009; Mori, 2022) toolbox in Python for visualization (each chord’s thickness is proportional to the weight of the functional connection; within-network edges are shown in the same color; between-community edge are shown in gray).

## Results

Subjects performed threat and reward trials at both low and high levels, in random order (Figure 1). The temporal window for analysis focused on the period preceding trial culmination as a virtual predator or a coin reward approached the bottom of the screen. The temporal window considered was defined to minimize potential contributions of transient responses at the onset of the trial (based on our previous work; McMenamin et al., 2014 and Murty et al., 2022).

First, we report skin conductance responses, which clearly ramped-up during the last seconds of the play period for both threat and reward conditions (Figure 2). In this figure and the ones below, the effect of arousal was defined as [(High_Threat + High_Reward) – (Low_Threat + Low_Reward)]; the effect of valence was defined as [(High_Threat + Low_Threat) – (High_Reward + Low_Reward)]. A 2 × 2 ANOVA detected a main effect of arousal but not valence (*F*(1,72) = 4.7/3.3, p=0.03/0.07, respectively); we did not detect an interaction effect (*F*(1,72)=0.4, p=0.53). Thus, in terms of autonomic responses captured via skin conductance, our manipulation was successful.

**Figure 2.**
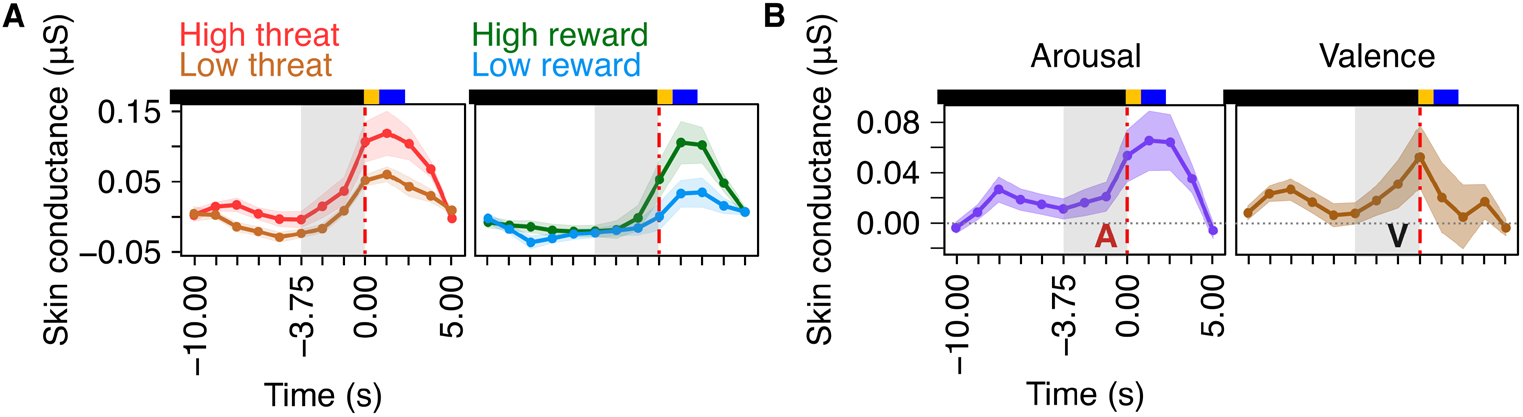
Skin conductance responses aligned to trial end (t = 0). The gray zone indicates the temporal window considered for analysis. (A) Responses for the four experimental conditions. (B) Temporal evolution of arousal and valence effects. The effect of arousal was statistically significant (red “A”), but no effect of valence was detected (black “V”). Colored bars above plots indicate trial timing: play period (black), indication period (yellow), and blank screen (blue; variable length).

Participants pressed buttons to attempt to escape from threat or try to catch the reward. We therefore analyzed two type of actions. Player movement was continuous as long as they pressed one of the two buttons (right/left movement). Total button-press times during the last 3.75 seconds before the play end were as follows (in seconds): high threat: 3.249; low threat: 3.252; high reward: 3.228; and low reward: 3.237. A 2 × 2 ANOVA did not detect main effects (valence: F(1,79)=0.125, p=0.724; arousal: F(1,79)=0.120, p=0.856) nor an interaction (F(1,79)=0.033, p=0.856). We also summarized the total number of motor actions (pressing or releasing a button). The average number of actions was as follows: high threat: 3.16, low threat: 3.0, high reward: 4.55, low reward: 4.317. Main effects were detected for both valence and arousal (F(1,79)=102.697/14.126, p=6.08×10^−16^/3.26×10^−4^, respectively); the interaction was not detected (F(1,79)=0.014, p=0.409).

### Functional MRI

Our first goal was to investigate responses in a set of regions of interest (ROIs) that is involved in aversive and/or appetitive processing based on the literature, including regions investigated in our previous studies (Table 1). A central aim was to test hypotheses related to the BST, amygdala, PAG, and ventral striatum. In addition, studies of sustained/uncertain threat have observed responses in the anterior insula but have not clearly separated the contributions of dorsal and ventral components; e.g. (Alvarez et al., 2011; Grupe et al., 2013; Hur et al., 2020). Here we test the prediction, based on our recent study (Murty et al., 2022), that the dorsal but not the ventral anterior insula would exhibit threat-related imminence responses. Likewise, we predicted that the anterior MCC but not the ACC would exhibit threat-related imminence-related responses. Figure 1B shows schematic responses based on increasing imminence-related responses starting at different times points prior to trial end. Statistical results for all ROIs are listed in Table 2.

**Table 2.**
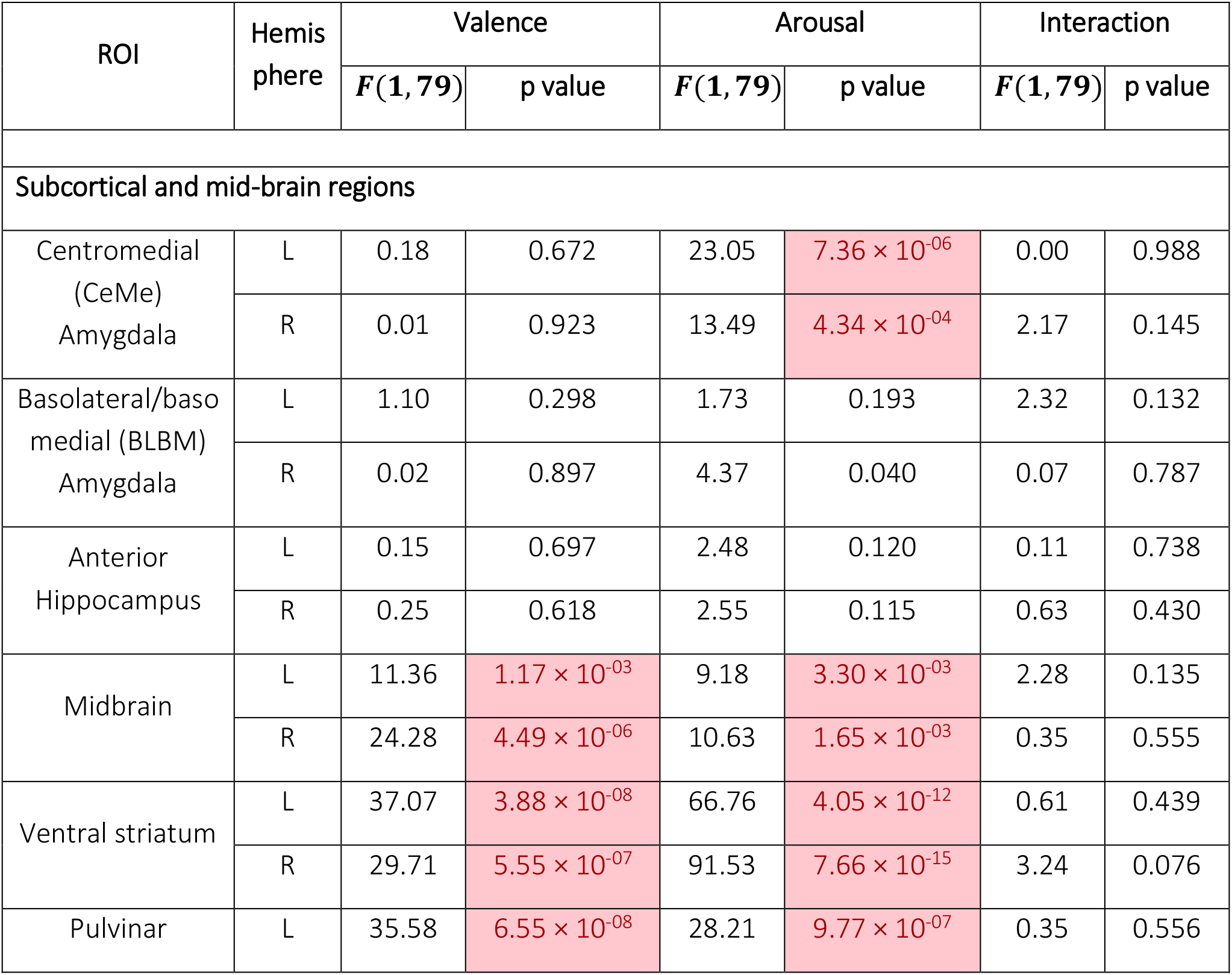
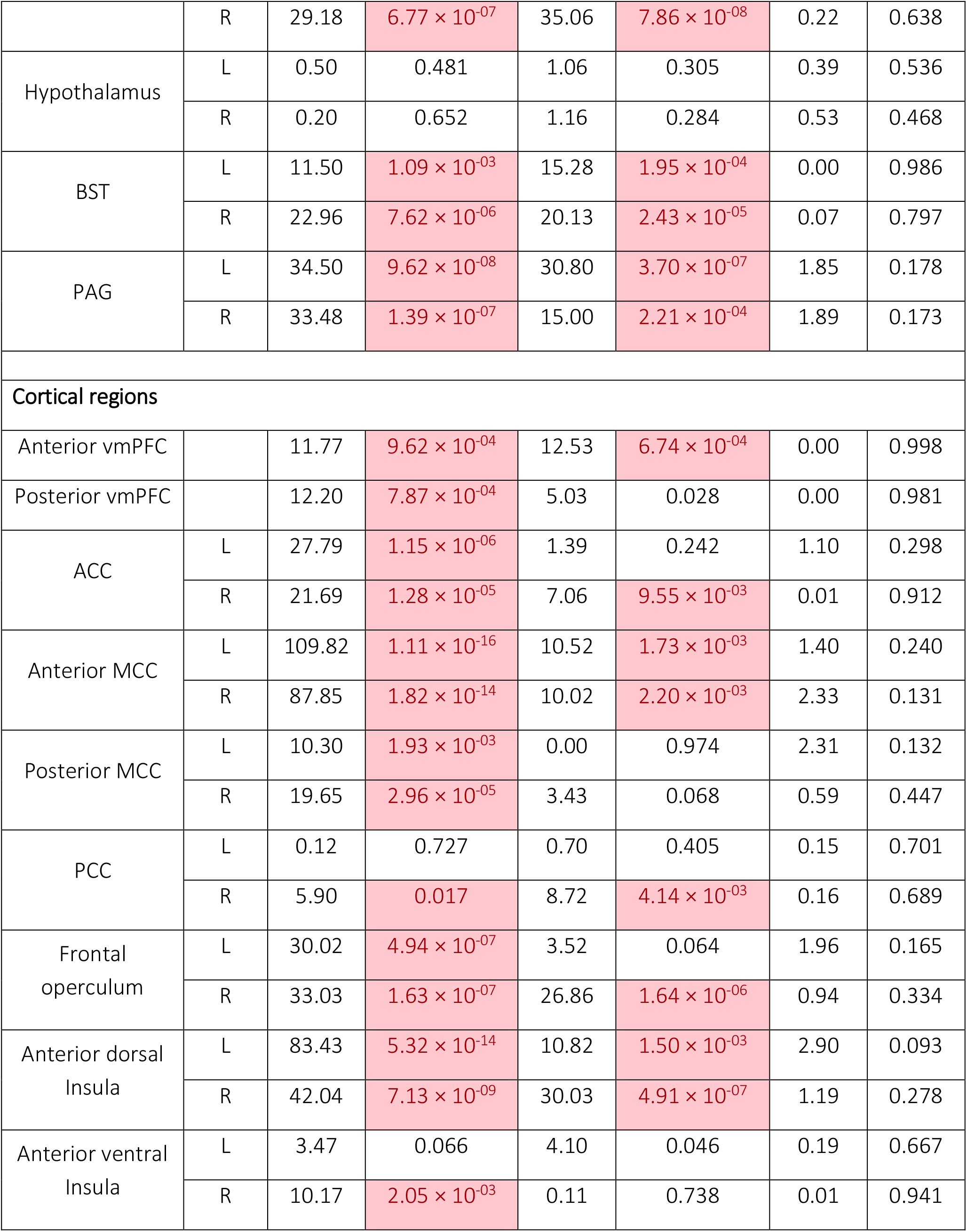
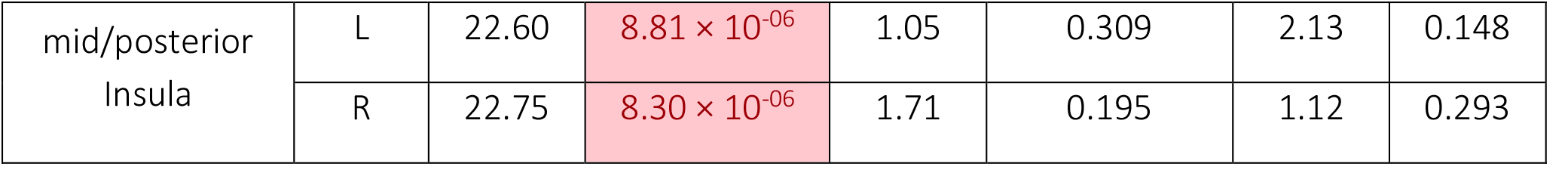
Statistical results for regions of interest: 2 Arousal x 2 Valence analysis of variance. Cells highlighted indicate that p-values survived False Discovery Rate correction.

The BST clearly exhibited ramping activity during the trial, whereas both amygdala ROIs displayed a qualitatively different response evolution (Figure 3). When considering the time course data in Figure 3 and subsequent ones, it is important to note that a response increase, such as the one observed at t = 0 for the BST, refers to events occurring ∼2 seconds earlier given the hemodynamic delay. Statistically, in the BST, the 2 × 2 ANOVA revealed main effects of arousal and valence; an interaction was not detected. In the centromedial amygdala and right basal amygdala (labeled BL/BM Amygdala in figures) an effect of arousal was detected but in the opposite direction: stronger responses were generated for low vs. high conditions. Both the response shape and the statistical results confirm our prediction of increased anticipatory responses during high threat trials in the BST, but not in the amygdala. Further, this difference across ROIs was confirmed by a 3-way ANOVA with ROI (BST, centromedial, and basolateral amygdala), valence, and arousal as factors, which revealed a significant interaction of arousal with ROI (*F*(1.6,126.3) = 27.4, p=3.3×10-9 for left and *F*(1.5,118.8) = 28.1, p=5.7×10-9 for right hemispheres). In addition, in the BST, the response evolution exhibited a very similar pattern during reward trials compared to threat trials; the main effect of arousal (without detecting an interaction) indicates that stronger responses were generated for high vs. low reward conditions. However, the BST was overall more strongly engaged by threat conditions as indicated by the main effect of valence. Like the BST, both the PAG and ventral striatum exhibited main effects of valence and arousal (no interaction was detected), and exhibited ramping activity during the trial end (Figure 4).

**Figure 3.**
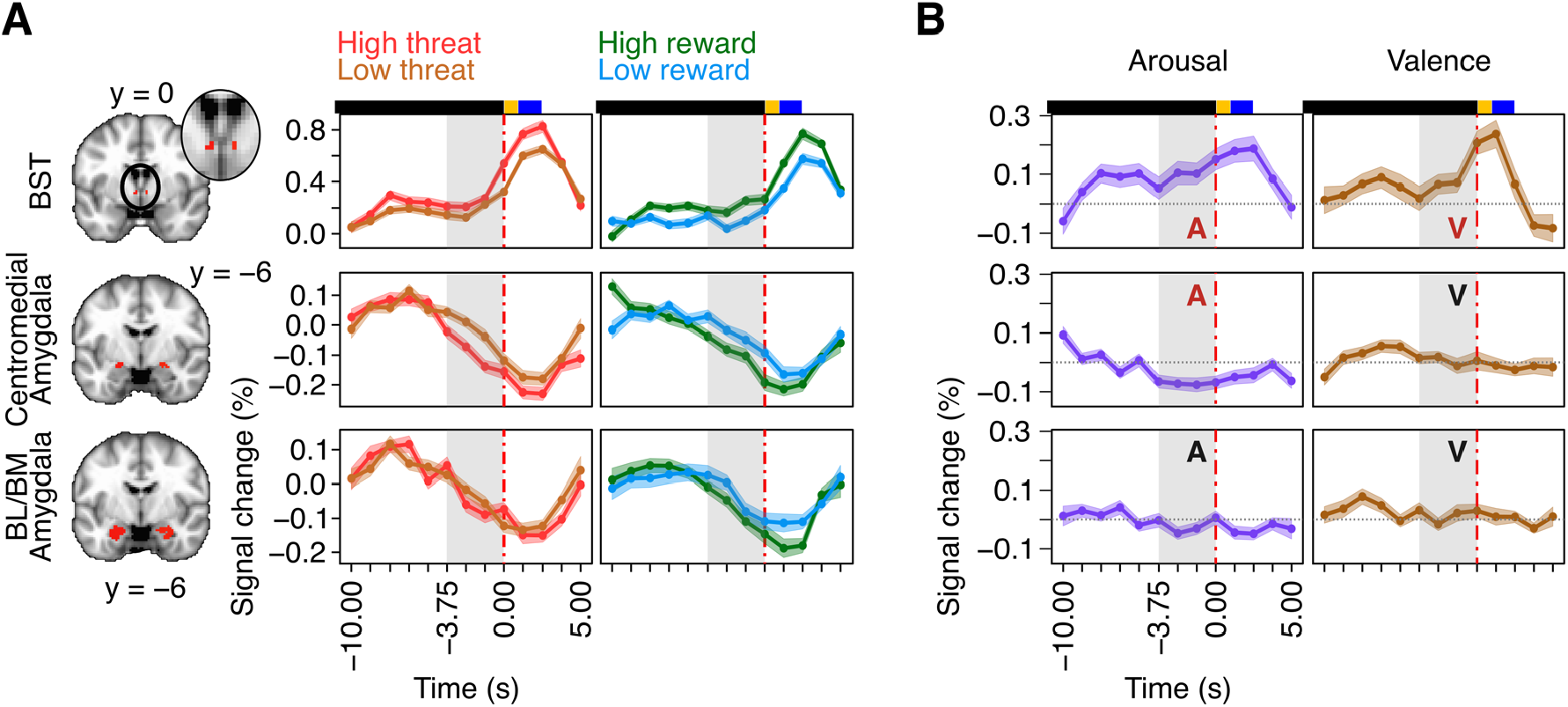
Estimated fMRI responses aligned to trial end (t = 0) for regions of interest (all in left hemisphere). The gray zone indicates the temporal window considered for analysis. (A) Responses for the four experimental conditions. (B) Temporal evolution of arousal and valence differential effects. Red letters “A” or “V” indicate that effects were detected at the significance level used; black letters indicate effects that were not detected. Colored bars above plots indicate trial timing: play period (black), indication period (yellow), and blank screen (blue; variable length). BL/BM: basolateral/basomedial.

**Figure 4.**
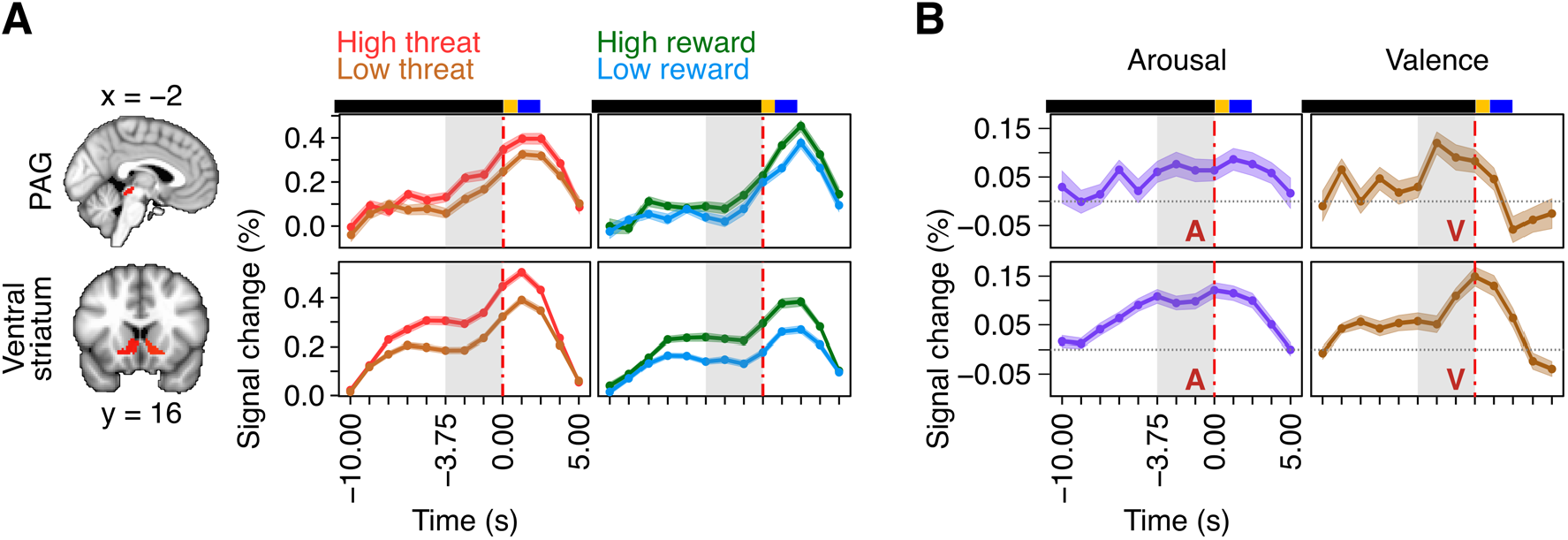
Estimated fMRI responses aligned to trial end (t = 0) for regions of interest (left hemisphere). Same format as in Figure 3. PAG: periaqueductal gray.

We characterized the responses in different sectors of the anterior insula (Figure 5). Whereas the dorsal ROI displayed a response pattern that followed that observed in the BST, PAG, and ventral striatum, the pattern in the ventral ROI was considerably different. Notably, the ventral ROI did not exhibit ramping responses but only an initial transient response at the onset of the trial. We statistically tested the ROI difference using a 3-way ANOVA with ROI (dorsal anterior insula and ventral anterior insula), valence, and arousal as factors, which revealed a significant interaction of ROI with arousal (*F*(1,79) = 21.3 / 30.3, p=1.5×10^−5^ / 4.5×10^−7^ for left / right hemispheres) as well as valence (*F*(1,79) = 22.6 / 16.2, p=8.7×10^−6^ / 1.3×10^−4^ for left / right hemispheres). For results comparing the anterior MCC and the ACC, see Figure 5. Whereas the anterior MCC displayed both an initial transient response and later ramping activity, the ACC only showed an initial transient response. A 3-way ANOVA with ROI (ACC and anterior MCC), valence, and arousal as factors revealed a significant interaction of ROI with arousal on the left side (*F*(1,79) = 12.3 / 0.5, p=7.3×10^−4^ / 0.4 for left / right hemispheres) as well as valence in both hemispheres (*F*(1,79) = 63.6 / 49.9, p=9.7×10^−12^ / 5.5×10^−10^ for left / right hemispheres), thus statistically confirming that imminence-related responses differed between the ACC and anterior MCC.

**Figure 5.**
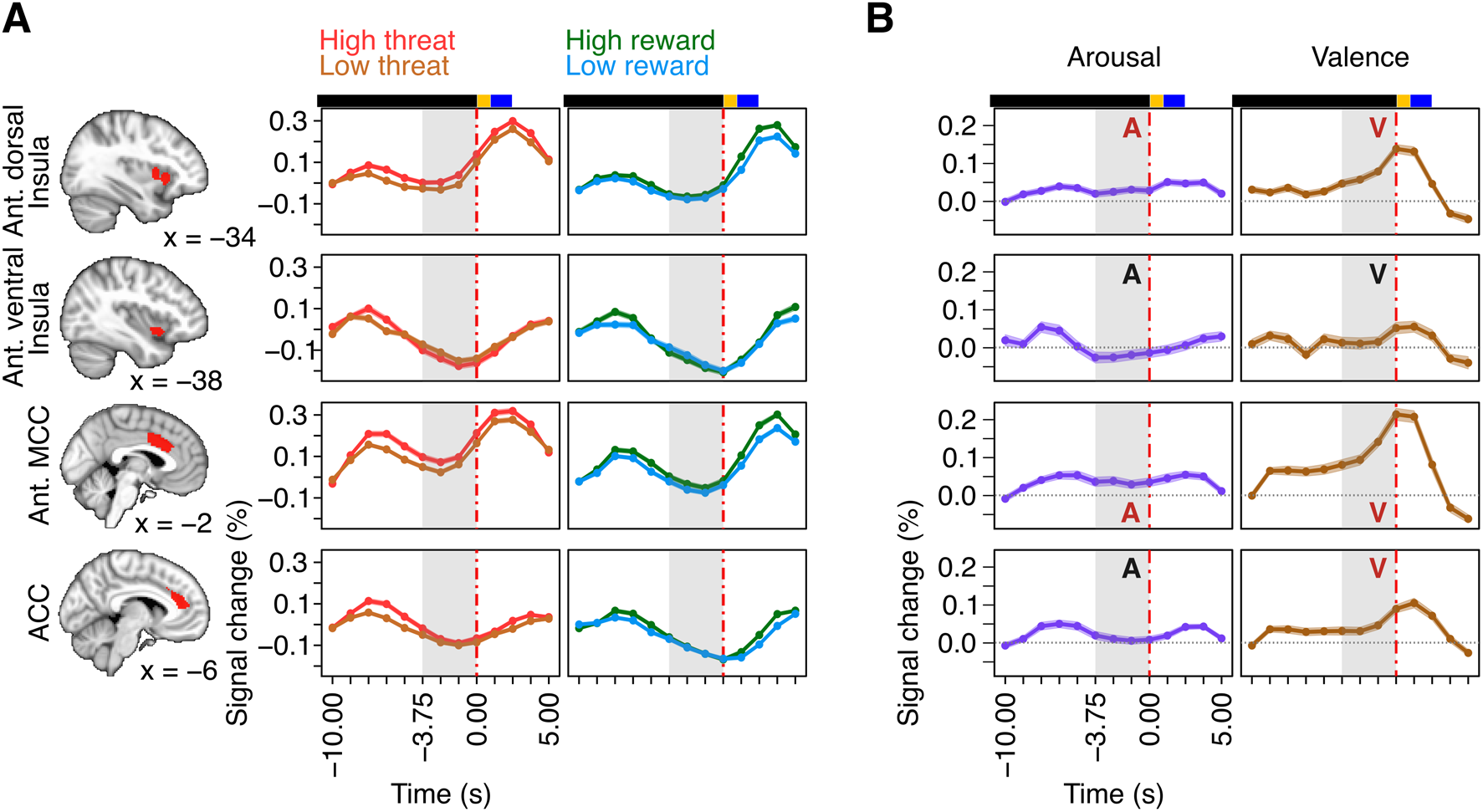
Estimated fMRI responses aligned to trial end (t = 0) for regions of interest. Same format as in Figure 3. ACC: anterior cingulate cortex; MCC: midcingulate cortex.

We also investigated the responses in two ROIs in the ventromedial PFC that have been implicated in safety-related processing. In these ROIs, responses tended to decrease along the entire anticipatory temporal window (Figure 6). Main effects of arousal and valence were observed in these regions (valence did not survive multiple comparisons correction for the posterior vmPFC), such that high arousal and threat conditions were associated with weaker signals.

**Figure 6.**
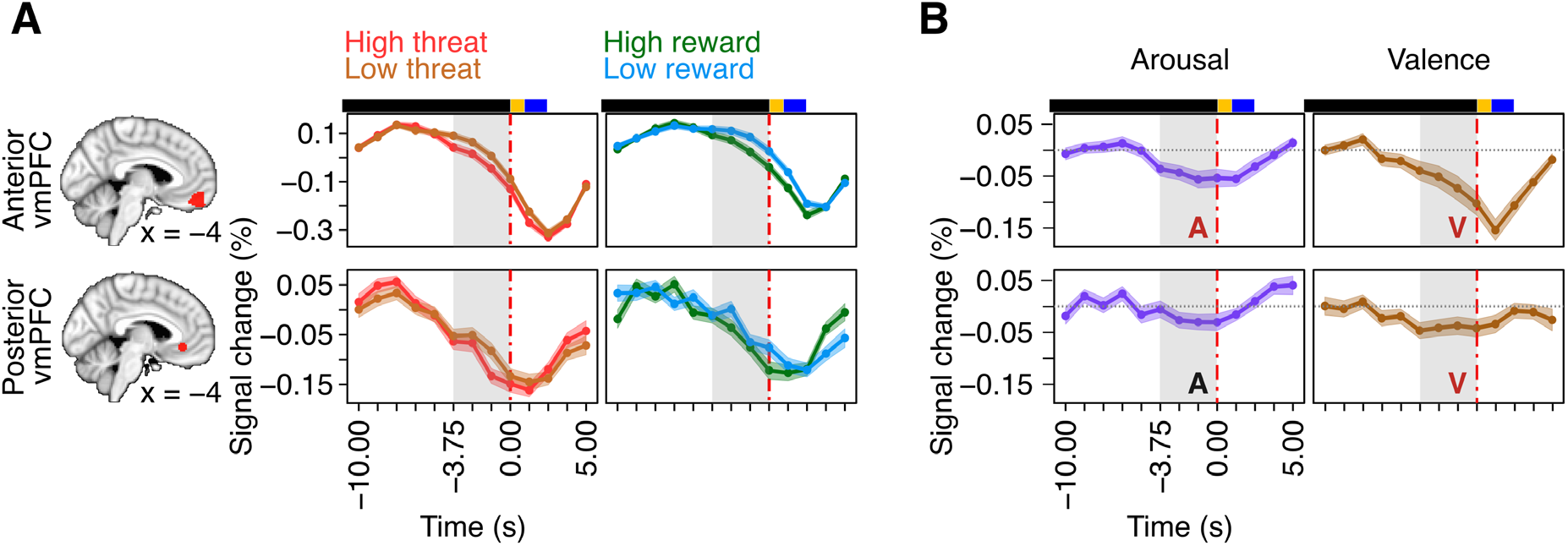
Estimated fMRI responses aligned to trial end (t = 0) for regions of interest (left hemisphere). Same format as in Figure 3. vmPFC: ventromedial prefrontal cortex.

#### Voxel-level analysis

We performed an additional voxelwise analysis to evaluate the impact of valence and arousal across the brain during the late trial phase. Both of these effects were detected quite widely, including cortical and subcortical sectors, in addition to brainstem and cerebellum, and overlapped spatially extensively (Figure 7). Figure 8 shows response patterns at some locations, including the fusiform gyrus, which is involved in face perception (icons used for threat trials included drawings of faces), and the frontal eye field, which is involved in spatial attention and eye movements.

**Figure 7.**
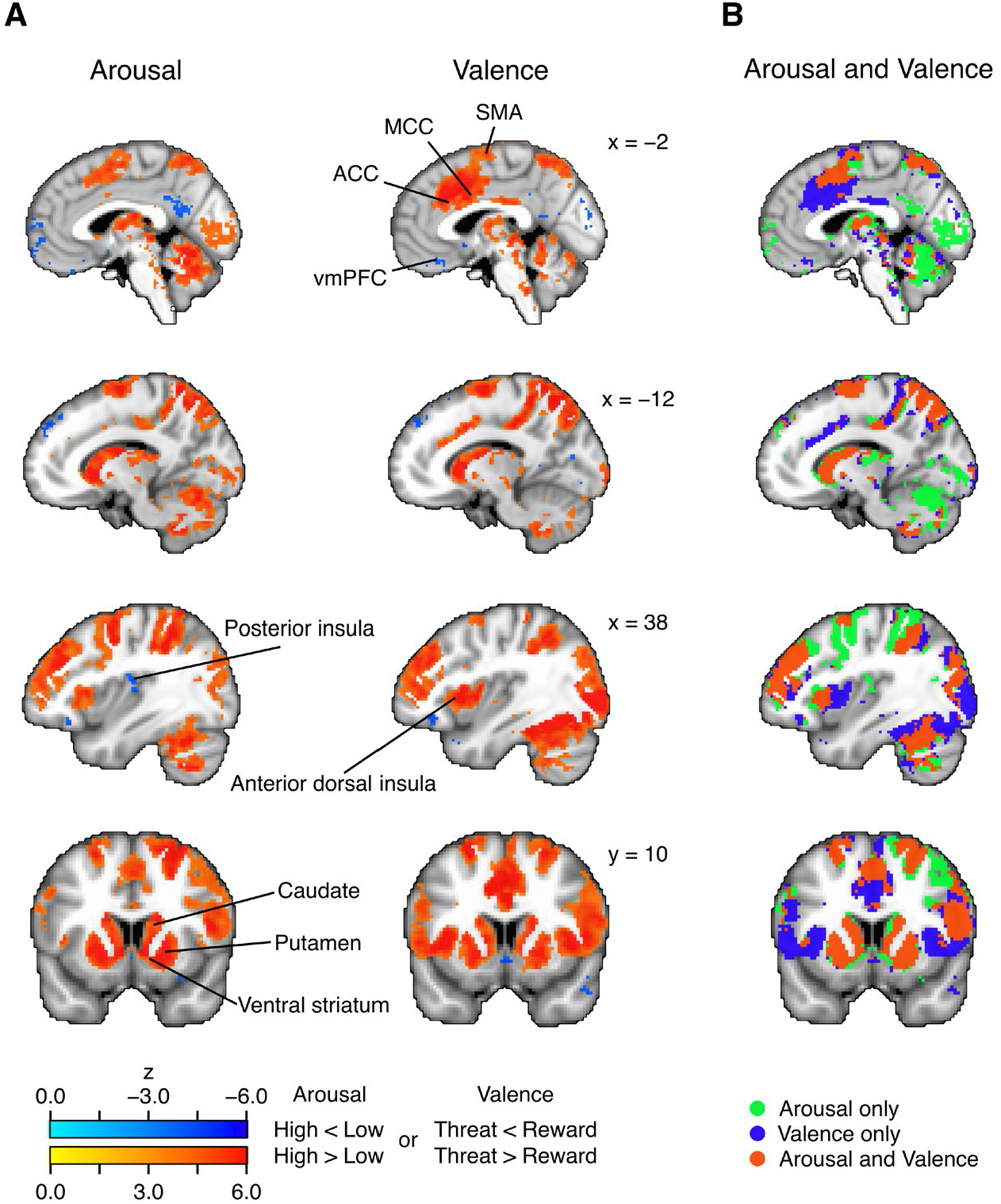
Voxelwise analysis. Effects of arousal and valence were thresholded using a false discovery rate of 0.001. The last column shows the same information as the arousal and valence maps but color coded to indicate where effects overlap spatially. ACC: anterior cingulate cortex; MCC: midcingulate cortex; SMA: supplementary motor area; vmPFC: ventromedial prefrontal cortex.

**Figure 8.**
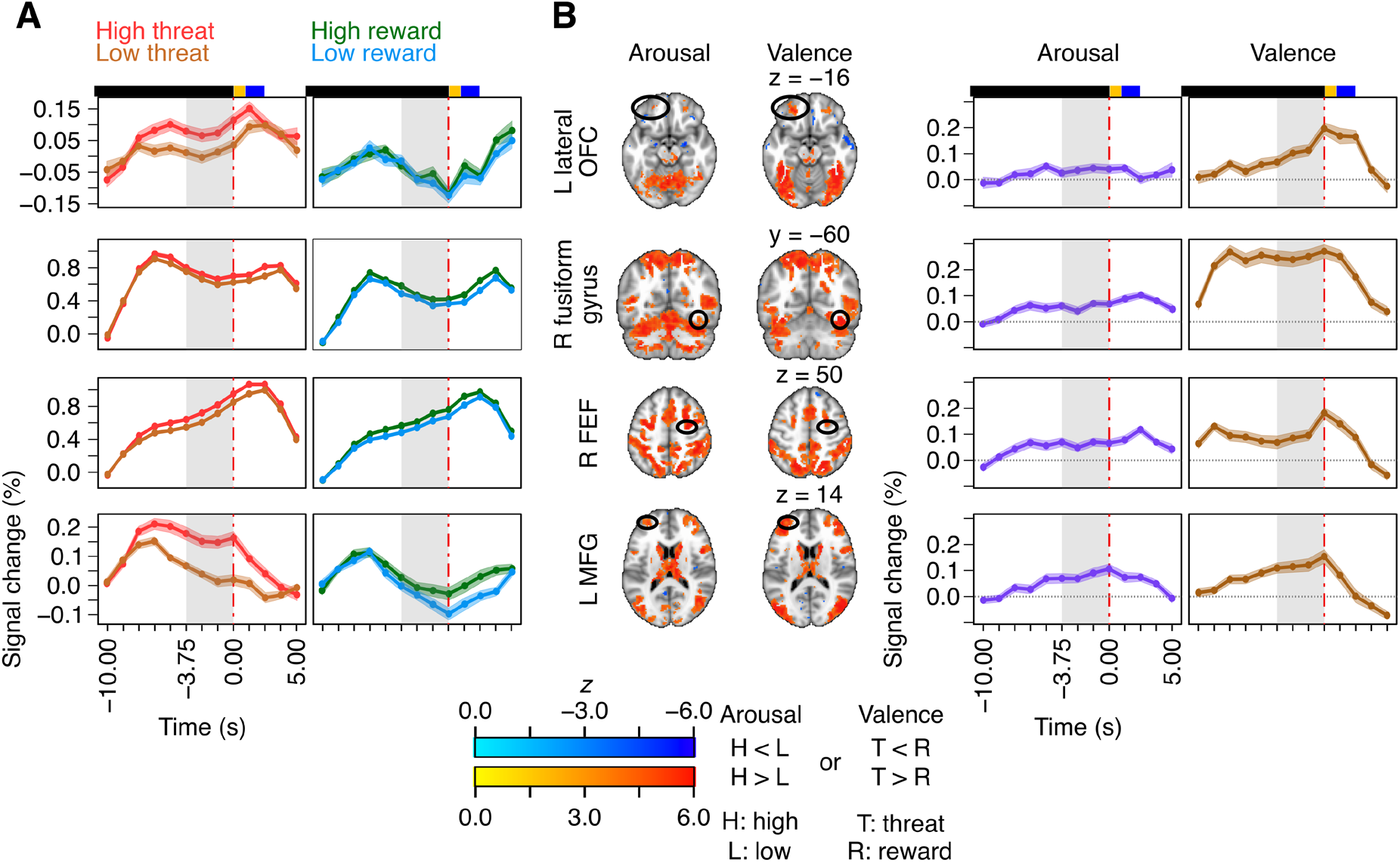
Estimated fMRI responses aligned to trial end (t = 0) for sets of voxels of the voxelwise analysis. Location of the voxels is indicated by black ellipses in the brain slices in (B). Effects were illustrated by averaging 7 neighboring voxels (center voxel plus 6 adjacent ones). The gray zone indicates the temporal window considered for analysis. (A) Responses for the four experimental conditions. (B) Temporal evolution of arousal and valence differential effects. Colored bars above plots indicate trial timing: play period (black), indication period (yellow), and blank screen (blue; variable length). FEF: frontal eye field; MFG: middle frontal gyrus; OFC: orbitofrontal cortex.

The voxelwise analysis also detected five clusters with valence by arousal interactions (Figure 9; we only considered clusters with at least 5 voxels surviving false discovery rate thresholding). Two of the clusters were in the so-called human MT+ complex that responds strongly to visual motion. In both clusters, the impact of arousal was larger for threat vs. reward conditions.

**Figure 9.**
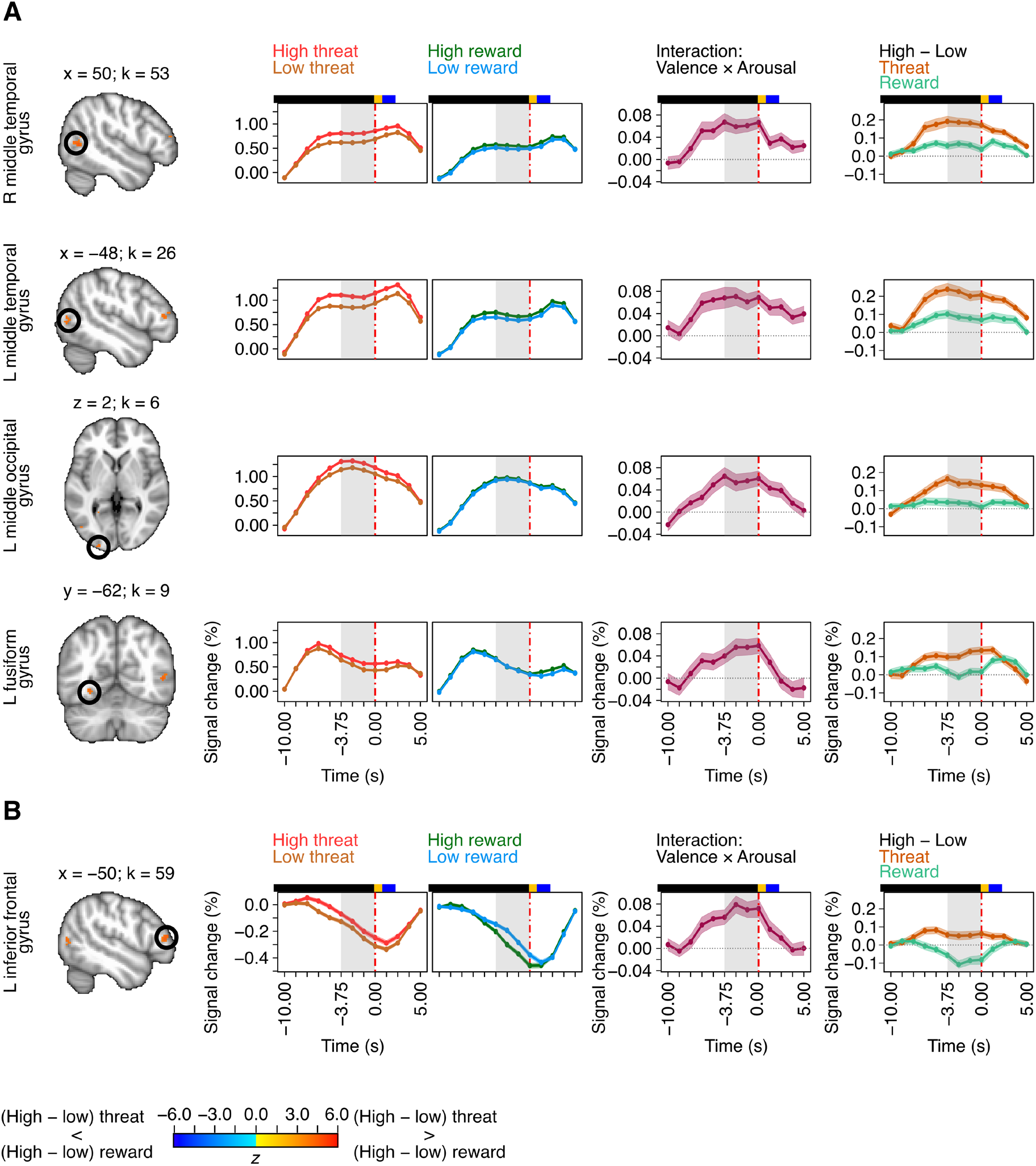
Valence by arousal interactions, voxelwise analysis. Effects are illustrated by averaging all voxels from clusters detected (cluster size indicated by *k*). Colored bars above plots indicate trial timing: play period (black), indication period (yellow), and blank screen (blue; variable length).

#### Exploratory network analysis

Based on a reviewer suggestion, we performed an exploratory network analysis to cluster brain regions into a set of networks (also called communities). We characterized networks for the high threat and high reward conditions, separately. In this manner, we were able to investigate how brain regions grouped into functional clusters based on trial-by-trial co-activation during the window of interest at the end of the trial (from -3.75 to 0 seconds) (Figure 10).

**Figure 10.**
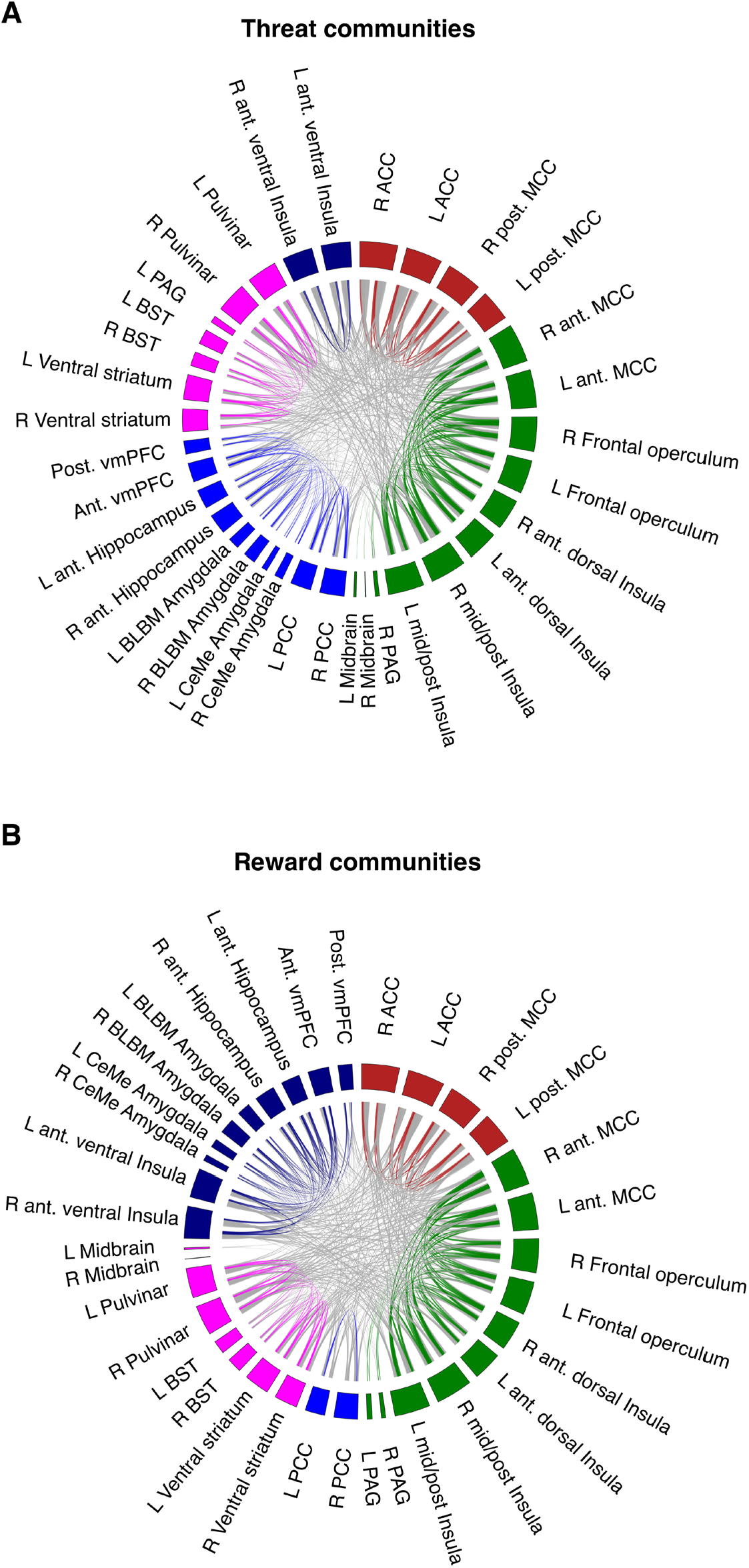
Functional networks. Regions of interest grouped into different communities (colors) based on imminence-related responses for high threat and high reward. The hypothalamus ROI was excluded because of poor SNR. ACC: anterior cingulate cortex; ant.: anterior; BLBM: basolateral/basomedial; CeMe: centromedial; L: left; MCC: midcingulate cortex; PAG: periaqueductal gray; PCC: posterior cingulate cortex. post.: posterior; R: right.

## Discussion

In the present study, we examined the effects of valence and arousal while subjects performed an avoidance/approach task. Whereas previous studies observed threat- and reward-related anticipatory responses in humans, our study was designed to test for the presence of increasing (or “ramping”) signals as a function of threat/reward imminence. Furthermore, our experiment allowed us to test the extent to which aversive- and appetitive-related processing recruit separate or overlapping territories. To uncover potential imminence-related responses, we estimated the temporal evolution of fMRI signals during the play period, which revealed such responses across multiple key brain regions involved in aversive and appetitive processing.

### Importance of studying threat and reward imminence dynamics

Several theoretical frameworks suggest that affective processes are inherently dynamic and/or involve multiple components that unfold temporally (see for example, Fanselow and Lester, 1988; Fanselow, 1994; Waugh et al., 2015; Sander et al., 2018; Mobbs et al., 2020). In addition, human behavioral studies have shown that dynamic stimuli (such as videos) enhance perceptual accuracy compared to static stimuli (such as pictures) across multiple paradigms, including recognition memory (Goldstein et al., 1982) and emotional face perception (Ambadar et al., 2005; Garrido-Vásquez et al., 2011; Dobs et al., 2018). In the context of nonhuman animal research, responses observed in areas like the amygdala have been shown to depend on task phase during dynamic paradigms. For example, Amir and colleagues (2015, 2019) showed that, when rats foraged under predatory threat, responses of distinct subpopulations of basolateral amygdala neurons were tied to particular task phases, such as trial initiation, foraging, or escape from predator. In another study, Gründemann et al., (2019) suggested that in their paradigm basal amygdala neurons did not signal “global anxiety” but predicted moment-to-moment switches between exploratory and non-exploratory defensive states. Thus, amygdala neurons did not appear to encode threat in ways that have been suggested when using static paradigms, but instead signaled distinct phases of “behavioral engagement” (Paré and Quirk, 2017). Taken together, these and related considerations motivated our investigation of the dynamic processing of threat and reward.

### Imminence-related responses in key regions of interest

We hypothesized that the BST and the PAG, two brain regions important for aversive processing (Davis et al., 2010; George et al., 2019), would exhibit imminence-related responses during threat, while the amygdala would not. In addition, we predicted that the dorsal anterior insula and the anterior MCC, but not the ventral anterior insula or the ACC, would behave in similar ways. Indeed, our findings confirmed these predictions. Furthermore, imminence-related responses were seen not only during avoidance conditions but also during reward approach, suggesting that these regions are sensitive to biologically significant conditions irrespective of valence.

#### Bed nucleus of the stria terminalis (BST)

The present results extend findings of the involvement of the BST during extended/uncertain threat and/or generating sustained responses during periods of anxious apprehension (Somerville et al., 2010, 2013; Alvarez et al., 2011; McMenamin et al., 2014; Meyer et al., 2019b; Hur et al., 2020; Murty et al., 2022) by demonstrating ramping activity with threat proximity. Surprisingly, such ramping signals were observed during reward trials, too. These results are consistent with the involvement of the BST in reward- and arousal-related processing in rodents (Jennings et al., 2013; Rodriguez-Romaguera et al., 2020), but previously not observed in the human literature, as far as we are aware.

#### Periaqueductal gray (PAG)

The PAG of the midbrain has been implicated in aversive and defensive reactions, and in mice PAG circuits control escape initiation (Bandler and Shipley, 1994; Branco and Redgrave, 2020). The virtual tarantula manipulation by Mobbs et al. (2010), where human participants were shown a prerecorded video of a spider moving toward or away from their feet, was effective in engaging the midbrain, including the PAG, when threat was proximal (although the activation was very extensive and thus difficult to localize). According to the threat imminence continuum framework (Fanselow and Lester, 1988), the PAG is a key node involved in circa-strike defensive behaviors (Fanselow, 1994). However, findings in the PAG have been somewhat inconsistent, possibly because of the challenges of scanning this midbrain region (which is adjacent to the cerebral aqueduct, and signal-to-noise ratio is compromised without additional procedures; see Methods). Whereas some studies found sustained activation to threat (Grupe et al., 2013; Hur et al., 2020), others only detected transient responses (Somerville et al., 2013), while some did not detect PAG responses (Somerville et al., 2010; Alvarez et al., 2011; McMenamin et al., 2014). Here, we observed ramping activity that started to rise 2.5 seconds before trial end (corresponding to events ∼4.5 seconds before trial end given the hemodynamic delay), consistent with the suggested role of this area. Like for the BST, we observed ramping activity during reward trials, too, albeit of weaker magnitude (note the robust valence effect), consistent with rodent work (Motta et al., 2017).

#### Cortical regions

Although some studies have suggested that the anterior insula in general is involved in affective processing, or that ventral sector is more strongly engaged in emotion-related processing whereas the dorsal in cognitive processes (Kurth et al., 2010; Uddin et al., 2017), here only the dorsal component generated imminence-related responses. Studies that reported increased activation to threat anticipation in the anterior insula (Alvarez et al., 2011; Grupe et al., 2013; Hur et al., 2020) did not explicitly subdivide the region into separate sectors, but inspection of the published maps suggests that the activation was mostly dorsal. In a recent study, we examined the two anterior insula sectors separately and found robust sustained threat-related responses in the dorsal but not the ventral anterior insula (Murty et al., 2022). Here, the dorsal anterior insula generated proximity responses to both aversive and appetitive conditions, suggesting that it is sensitive to motivationally significant information irrespective of valence (Snellenberg and Wager, 2009).

We observed a related pattern of results in the context of different cingulate territories. Specifically, the anterior MCC exhibited anticipatory responses for both valences, but not the ACC, consistent with results by Grupe et al. (2013) and Hur et al. (2020) (but see Alvarez et al., 2011 and McMenamin et al., 2014). The anterior MCC is a motivational hub believed to play a major role in threat assessment and/or appraisal (Shackman et al., 2011; Kohn et al., 2014; Vogt, 2016; Langner et al., 2018). We previously suggested that the anterior MCC is a site of interaction of negative valence and motor-related signals (Pereira et al., 2010; Lima Portugal et al., 2020). However, here we also observed evidence for anticipatory responses during reward trials, supporting the notion that several brain regions are sensitive to motivationally significant information irrespective of valence. The voxel-level maps provided additional information of valence and arousal effects in medial prefrontal cortex. We observed main effects of both valence and arousal in more dorsal aspects of the dorsomedial PFC, while the effect of arousal was not detected in the relatively more ventral anterior MCC.

#### Ventral striatum

Based on prior literature (for review, see Knutson and Greer, 2008), we anticipated that the ventral striatum would exhibit ramping activity during reward trials. Our results revealed that signals increased with potential reward proximity, at least for the last few seconds of the trial, in a manner resembling that in rodents (Howe et al., 2013). In addition, we tested the hypothesis that imminence-related responses would be generated during aversive processing, too, consistent with findings demonstrating a role of the ventral striatum during escape behaviors (LeDoux et al., 2017; Ray et al., 2022). Indeed, our results uncovered the involvement of this region during the last seconds of threat trials, consistent with a role in escape behaviors, in contrast to previous studies that did not permit escape from threat (McMenamin et al., 2014; Murty et al., 2022).

### Role of amygdala during active aversive and appetitive conditions

Although a considerable literature has documented the involvement of the human amygdala in threat-related processing, this area appears to be more tied to the processing of temporally precise threat (Davis et al., 2010). However, it is also important to note that at a large meta-analyses did not detect amygdala responses during fear conditioning in humans (Fullana et al., 2016; but see Wen et al., 2022); another meta-analysis did not detect reliable amygdala involvement during uncertain threat (Chavanne and Robinson, 2021). Furthermore, several studies have observed decreases in amygdala responses during particular threat paradigms, including from out lab (Pruessner et al., 2008; Choi et al., 2012; McMenamin et al., 2014; Visser et al., 2021; Murty et al., 2022). In the present study, both the basal and the centromedial amygdala ROIs showed no evidence of temporally increasing responses during threat trials. In fact, not only responses decreased during the trial, but high threat signal decreased more strongly. Our results thus conflict with the view that the human amygdala is engaged during extended periods of threat (Hur et al., 2020).

### Ventromedial PFC

Previous studies have suggested that the ventromedial PFC signals relative safety (Schiller et al., 2008; Eisenberger et al., 2011) and/or that the region evokes stronger responses during less versus more threatening conditions (Murty et al., 2022). In the present study, the temporal evolution of signals in the ventrolateral PFC regions was qualitatively different from those observed in regions such as the BST and dorsal anterior insula, for example. Instead, responses decreased during the trial period, a pattern that was also observed in both amygdala regions. In the anterior ventromedial PFC, we found stronger responses to low versus high threat, consistent with the prior literature. However, a similar finding was detected during reward trials, a finding that runs counter the idea that the ventromedial PFC signals different levels of threat.

### Voxelwise analysis

The voxelwise analysis revealed that both the effects of valence and arousal were quite spatially extensive, spanning cortical and subcortical sectors, in addition to brainstem and cerebellum. Notably, our study uncovered a considerable contribution of the main effect of arousal (without detecting an interaction; but see below) across many cortical, subcortical, brainstem, and cerebellum regions, indicating that most regions engaged by imminent threat were also engaged by imminent reward.

Surprisingly, in the ROI analysis, we did not detect an arousal by valence interaction. In the voxelwise analysis, we detected five sites with interaction patterns. We only observed one cluster with a cross-over type of interaction, where the effect of arousal was in one direction for threat and the opposite for reward (left inferior frontal gyrus). But given that this cluster did not show increasing imminence-related responses, it is harder to interpret this response pattern. Furthermore, a few other clusters without interaction effects displayed qualitatively different responses during threat and reward trials. This was the case in the lateral orbitofrontal cortex, which generated more sizeable imminence-related responses during threat trials. In addition, the middle frontal gyrus showed sustained responses throughout most of the trial period, but only more robustly for high threat trials. Finally, our study also detected an interaction pattern in the human MT+ complex, a region that is strongly driven by visual motion (Culham et al., 2001). In two clusters, arousal had a greater impact during threat vs. reward trials.

### Attention and motor responses

Many of the regions investigated showed qualitatively similar temporal response evolution for both threat and reward conditions. This finding was surprising to us, especially in the subcortical regions we targeted (e.g., BST and PAG). Do the similar response patterns reflect general contributions of attention? It is reasonable to assume that participants paid increased attention to conditions involving high vs. low arousal. In this context, it is instructive to consider the responses in the frontal eye field, a region strongly engaged by conditions involving spatial attention and eye movements (Grosbras and Paus, 2002; Muggleton et al., 2003). The response evolution in this region was considerably different from that observed in our key ROIs, and ramped up very early and not only during the last few seconds of the trial; for example, taking the hemodynamic response into account, BST signals started to ramp up around 3.75 seconds prior to trial end. It is also instructive to consider responses in the fusiform gyrus, a region that is engaged by face stimuli (recall that threat icons contained faces) and modulated by spatial attention (Wojciulik et al., 1998). Although the fusiform gyrus exhibited arousal and valence effects, we did not observe a pattern of ramping activity with threat/reward proximity.

Thus, whereas it is possible that attentional processing contributed to some of the results we observed, it is unlike to explain key properties of the temporal evolution of signals during the analysis window at the end of the trial. More broadly, periods of threat and reward imminence likely engage multiple mental processes, including those traditionally described as attentional, motivational, emotional, and action related. Whereas there is value in attempting to isolate some of these processes (e.g., “purely emotion related”), in more natural settings, they are jointly engaged. In fact, as argued elsewhere, we believe that trying to disentangle them can be counterproductive (Pessoa, 2022; Pessoa et al., 2022). As researchers start embracing increasingly dynamic and naturalistic experimental paradigms, it will be critical to consider the inherent intertwining of diverse mental domains (“attention” and “emotion”, say).

How about contributions of motor actions? Total time pressing buttons was very similar across all conditions, so this factor probably did not contribute to differential responses. However, we did detect effects of arousal and valence on the total number of actions, with more of them occurring during high vs. low conditions. It is thus possible that motor actions contributed to the responses observed in some of the regions. However, note that the largest number of motor actions was generated during high reward trials, but fMRI responses during reward trials were not greater than during threat trials in general. Thus, whereas motor actions might have contributed to the effects of arousal, in particular in regions more sensitive to motor-related signals, they do not explain the results in our key regions. For example, the anterior MCC is particular tuned to actions, but responses in this region were largest for threat relative to reward trials.

### Exploratory network analysis

The investigation of activation identified sets of regions with similar responses. To further explore the possibility that these regions formed functional clusters, we performed graph analysis to identify potential networks based on trial-by-trial co-fluctuations during the temporal window of interest. Notably, during high threat, a network involving the pulvinar, PAG, BST, and ventral striatum was identified. Based on their activation profiles and their response co-fluctuation, we propose that they form an important network engaged by the processing of imminent/proximal threat. Importantly, amygdala ROIs and the anterior hippocampus did not group with this network and were part of a cluster that included both ventromedial PFC regions and the posterior cingulate cortex. These results further strengthen the notion that the amygdala and anterior hippocampus are not strongly involved in temporally extended threat in humans. During reward trials, similar networks were observed, with some notable differences. For example, during reward trials, the PAG grouped with another network, whereas the midbrain grouped with the pulvinar, BST, and ventral striatum. Taken together, our analysis provides an initial working model for imminent-related processing in the context of threat and reward, one that suggests that the associated networks exhibit considerable overlap.

## Conclusions

We uncovered “ramping” imminence-related responses during threat trials in multiple brain regions, including the BST and the PAG (but not the amygdala). Whereas the ventral striatum generated anticipatory responses in the proximity of reward as expected, it also exhibited threat-related imminence responses. In fact, across many brain regions we observed a main effect of arousal ([high threat + high reward] > [low threat + low reward]). In other words, we uncovered extensive temporally-evolving, imminence-related processing in both the aversive and appetitive domain, suggesting that distributed brain circuits are dynamically engaged during the processing of biologically relevant information irrespective of valence, findings further supported by exploratory network analysis.

## Acknowledgments

This research was supported by the National Institute of Mental Health (R01 MH071589 and R01 MH112517). We thank Joyneel Misra for assistance with coding the experiment, and Navot Naor for early work leading to the development of the paradigm studied here.

